# Language models learn to represent antigenic properties of human influenza A(H3) virus

**DOI:** 10.1101/2025.01.17.633534

**Authors:** Francesco Durazzi, Marion P. G. Koopmans, Ron A. M. Fouchier, Daniel Remondini

## Abstract

Given that influenza vaccine effectiveness depends on a good antigenic match between the vaccine and circulating viruses, it is important to assess the antigenic properties of newly emerging variants continuously. With the increasing application of real-time pathogen genomic surveillance, a key question is if antigenic properties can reliably be predicted from influenza virus genomic information. Based on validated linked datasets of influenza virus genomic and wet lab experimental results, *in silico* models may be of use to learn to predict immune escape of variants of interest starting from the protein sequence only. In this study, we compared several machine-learning methods to reconstruct antigenic map coordinates for HA1 protein sequences of influenza A(H3N2) virus, to rank substitutions responsible for major antigenic changes, and to recognize variants with novel antigenic properties that may warrant future vaccine updates. Methods based on deep learning language models (BiLSTM and ProtBERT) and more classical approaches based solely on genetic distances and physicochemical properties of amino acid sequences had comparable performances over the coarser features of the map, but the first two performed better over fine-grained features like single amino acid-driven antigenic change and *in silico* deep mutational scanning experiments to rank the substitutions with the largest impact on antigenic properties. Given that the best performing model that produces protein embeddings is agnostic to the specific pathogen, the presented approach may be applicable to other pathogens.

## Introduction

Thanks to the advancements in next-generation sequencing and the increase of computational power, genomic surveillance is emerging as a first-line tool to monitor infectious disease outbreaks, inform public action in real-time, and inform and evaluate intervention strategies. The availability of whole genome sequencing data and novel phylogeny algorithms allows the rapid identification of new variants, which may be associated with new functional properties such as different infectiousness, virulence, resistance to therapeutics and escape from natural or vaccine-induced immunity^1,2^.

Traditional phylogeny allows reconstructing the history of virus emergence and evolution, including the identification of transmission chains, and super-spreading events^3^. Prediction of functional traits, however, is more challenging, for instance when linking mutations with antigenic properties or virulence. Therefore, we explored the use of machine learning approaches to recover this information from protein sequence alone, complementing and supporting experimental measurements (e.g. prioritizing variants in inhibition assays).

The discrete and symbolic (i.e. non-metric) nature of protein sequences can be tackled through Natural Language Processing (NLP) methods^4,5^, such as Language Models (LM) that transform amino acid (AA) sequences into fixed-length vectors that optimally encode the “linguistic” content of the sequence. Deep Learning models commonly used for this task are recursive networks, such as Bi-directional long-short-term memory neural networks (Bi-LSTM)^6^, or attention-based transformers (i.e. BERT ^7^), which also allow faster training and easier interpretability.

Recently, Hie et al.^8^ trained a BiLSTM language model on approximately 50,000 influenza A(H1) virus hemagglutinin (HA) sequences and showed that a combination of distance in the embedding space (“semantic” change from a reference) and probability of specific mutations (“grammaticality” of the mutation) was predictive of immune escape properties of viral isolates. The capability to predict antigenic properties of influenza viruses directly from protein sequences would be of great help for the Global Influenza Surveillance and Response system (GISRS)^9^ that is in place for vaccine strain selection to reduce the impact of this disease, which still puts a significant burden on public health in terms of morbidity, deaths and associated costs^10,11^. Antibodies against HA can provide protective immunity to influenza virus infection or disease, and for this reason this protein is a primary component of vaccines^12^. Antigenic differences between vaccine strains and circulating viruses can cause reduced vaccine effectiveness and have therefore been routinely evaluated in surveillance- and vaccine development programs over the years. To this end, hemagglutination inhibition (HI) assays, which measure the ability of antisera to block the virus-mediated agglutination of red blood cells, are routinely used as an appropriate surrogate test for the more time-consuming virus neutralization assays^13^. For a quantitative interpretation and visualization of HI data, antigens and antisera can be represented in an antigenic map, such that the distance between antigens and antisera in the map are inversely related to the HI titers^14^. This method was initially applied to a dataset of 273 influenza A(H3N2) virus isolates in 2004^12^, which was extended to 279 viruses in 2011^15^.

HA1 sequence data were generated for the same dataset. The higher-level structure of the antigenic map was found to be punctuated rather than gradual with 13 clusters of antigenically related viruses appearing in chronological order, reflecting selection of variants with increased fitness in the background of population immunity built up against previously circulating variants. The AA substitutions responsible for the major antigenic changes between viruses from one cluster and the next have been mapped using reverse genetics^16^. Surprisingly, relatively few AA substitutions were found to cause large differences in antigenic properties, while the majority of genetic changes had little or no antigenic effect. As a consequence, ordinary measures of (phylo)genetic distance do not capture antigenic evolution well.

Here we explored the possibility to predict antigenic map coordinates directly from protein sequences without the need of HI experiments, which would be of great help in the current era of sequence-based surveillance. In previous studies, machine learning was employed to predict antigenic distances starting from genetic differences between pairs of viruses, computed from substitution matrices^17,18^ and additional expert-curated features based on known relevant positions, e.g. around the receptor-binding domain^19^. Here, instead of trying to reconstruct the antigenic distance matrix, we represented the HA sequence of each isolate as a numerical vector (obtained with different methods based on LMs, genetic or physicochemical properties) and directly predicted its antigenic map coordinates through regression.

We compared four methods to represent protein sequences as numerical vectors: two of which were obtained through Deep Learning language models (one of which is trained specifically for influenza virus HA and the other is more agnostic regarding the specific protein) and one was based on a signature of physicochemical properties at the single AA level derived from AAindex^20^. We compared these methods against a representation based solely on Hamming distance between AA sequences as a benchmark. We compared a) the performance in predicting antigenic map coordinates, b) the sensitivity to single AA substitutions driving antigenic change and c) the generalizability of antigenic map prediction to unseen antigenic clusters.

#### How LM protein sequence embeddings work

Language models are trained to predict in an unsupervised way “tokens” within a “sentence” (e.g. AAs within a protein sequence or words within a text sentence) using the rest of the sentence as a context. Different LMs may have different architectures (e.g. recursive networks like BiLSTM or transformer networks like BERT), but in all cases the output of one of the internal artificial neuron layers is used as a vector representation of the input sentence, called *embedding*. It is assumed that the higher the accuracy of sentence reconstruction, the more informative its embedded representation. The embeddings can then be used to recover additional information, like antigenic properties of the sequence, through classification or regression algorithms.

## Materials & Methods

### Data

Antigenic map coordinates (or AM space) were available for 279 antigen samples, corresponding to 209 unique protein sequences^12,15^. We averaged the AM coordinates over the samples with identical HA protein sequence, considered as technical replicates. The mean standard deviation of antigenic distance for samples sharing the same HA protein sequence was 0.46 a.u. on the x-axis and 0.59 a.u. on the y-axis, which is below the experimental error estimated at 0.83 a.u.^12^. The target antigenic map has been obtained through multi-dimensional scaling (MDS) of the hemagglutinin inhibition (HI) titers generated with a set of influenza A (H3N2) viruses and a panel of ferret antisera against these viruses, as described in ^12,15^. MDS generates a vector space of chosen dimensionality in which distances between antigens and sera are inversely related to the HI titers. In ^12^ the MDS space was calculated to require dimensionality d=2, which we refer to as MDS1 and MDS2 in the figures.

### Genetic distance embeddings

We computed the pairwise Hamming distance between aligned protein sequences by counting the number of different AAs. We then applied MDS on the distance matrix to obtain a vector embedding for each virus. To perform the antigenic map regression in similar conditions for the different embedding methods (e.g. in terms of Vapnik-Chervonenkis dimension^21^), we mapped the Hamming distances into a 1024-dimensional vector space, analogously to LM embeddings. This was done in order to minimize the risk of overfitting by one method due solely to its embedding dimensionality.

### Physicochemical signature

We represented each AA of the HA sequences with 3 physicochemical properties^16^, namely charge (C), volume (V), and hydropathy index (H) (extracted from AAindex^20^, a database of >600 chemo-physical properties for each AAs). We thus embedded each HA sequence as a 3×329=987-dimensional vector. The physicochemical signature is the only representation we tested that has a direct relationship between vector elements and AA positions along the sequence (i.e. the first 3 vector components refer to C,V,H values for the first AAs of the HA sequence, and so on). We tested a signature of similar size composed by the first 3 Principal Components of the whole set of AAindex properties, but the results were comparable, thus we considered the C,V,H signature because it has a clearer interpretation.

### BiLSTM embeddings

We used a Bi-directional Long Short-Term Memory neural network (BiLSTM), trained on a dataset of 45K influenza virus hemagglutinin sequences of multiple subtypes and collected from multiple animal hosts as in ^8^. We started from the work as in ^8^, from which we took the Language Model and the full training data, and then retrained and tested it on our servers using the Python package Tensorflow. To maximally avoid overfitting in the antigenic map regression performed later, we removed from the original BiLSTM training set all the HA sequences matching the 279 sequences used in the antigenic map dataset. In Language Models, the network is trained to reconstruct a protein sequence after masking one AA at a time along the whole sequence. The output of the second-last layer of the network was used as embedding of the protein sequence.

### ProtBERT embedding

ProtBERT is a BERT (Bidirectional Encoder Representations from Transformers) language model trained on two large curated protein datasets: UniREF (www.uniprot.org/uniref) and BFD (bfd.mmseqs.com), containing up to 393 billion AAs from 2.5 billion protein sequences sampled along the full tree of life^22^. Training of this model exceeds the computational capabilities of our laboratory, thus we used the pre-trained model as provided by the authors (available through the Python package in Huggingface website). Both ProtBERT and BiLSTM embeddings have been computed on a Nvidia A100 GPU with 80GB VRAM, requiring up to 500GB RAM and 24 hours (for BiLSTM, which is slower).

### Ridge regression (RR)

We trained four Ridge regression models to predict the bi-dimensional antigenic map coordinates: one RR for each of the previously described protein embeddings. The independent variables were the embedding vectors, which we standardized before regression. The target variables were the 2 AM coordinates, spanning 17 antigenic units (a.u.) for MDS1 and 33 a.u. for MDS2. One a.u. of antigenic distance between antigen and antiserum corresponds to a 2-fold dilution of antiserum in the HI assay. A validation set was generated by randomly sampling 26 sequences from the AM set, ensuring that at least 1 sequence for each antigenic cluster is present in the test set. The Ridge regression was trained on the remaining 183 sequences with Leave One Out Cross-Validation (LOO-CV), by averaging the coefficients of the 183 (=209-26) LOO regressors. We also tested nonlinear regression methods (e.g. Feedforward Neural Networks) but no improvement in the performance was observed thus we kept the simplest model.

### Leave-the-Future-Out (LFO)

To test the generalizability of the approach to future antigenic clusters, totally unseen during RR training, we removed from the RR training set the most recent influenza virus antigenic cluster (PE09). We then predicted the positions of antigens held out with RR. We only aimed to verify whether sequences belonging to PE09 were predicted to be placed outside the previous cluster (CA04), given that the exact directionality of placement can not be determined without antisera against the newly emerging variant and titrations against older ones. We considered a new sample to be outside the CA04 cluster either according to a 95% confidence interval estimated with a 2D Gaussian probabilistic fit of the CA04 cluster, or if it was predicted to be 2 a.u. distant from the center of the CA04 cluster.

## Results

### Antigenic map prediction performance

Before regression from protein embeddings to AM coordinates, we evaluated the accuracy of the language models in reconstructing the AA HA1 protein sequences of the AM set. We tested 3 different architectures of the hidden LSTM layer (embedding size = 128, 512, and 1024), and with 1024 dimensions we obtained better results both in language model accuracy and in antigenic map prediction (see Supplementary Table S1). For BiLSTM, the reconstruction performance was 0.98 +-0.02, which means that 98% of AAs were correctly predicted in the AM dataset. For ProtBERT it was 0.99 +-0.01, compatible with BiLSTM performance.

We next performed ridge regression on AM coordinates, starting from the 4 embeddings introduced above. In Table 1 we show the prediction errors for leave-one-out predictions. The errors were computed as the Mean Average Error (MAE) between true and predicted antigenic map coordinates, shown both for the training and the validation set (see Methods).

**Table 1.**
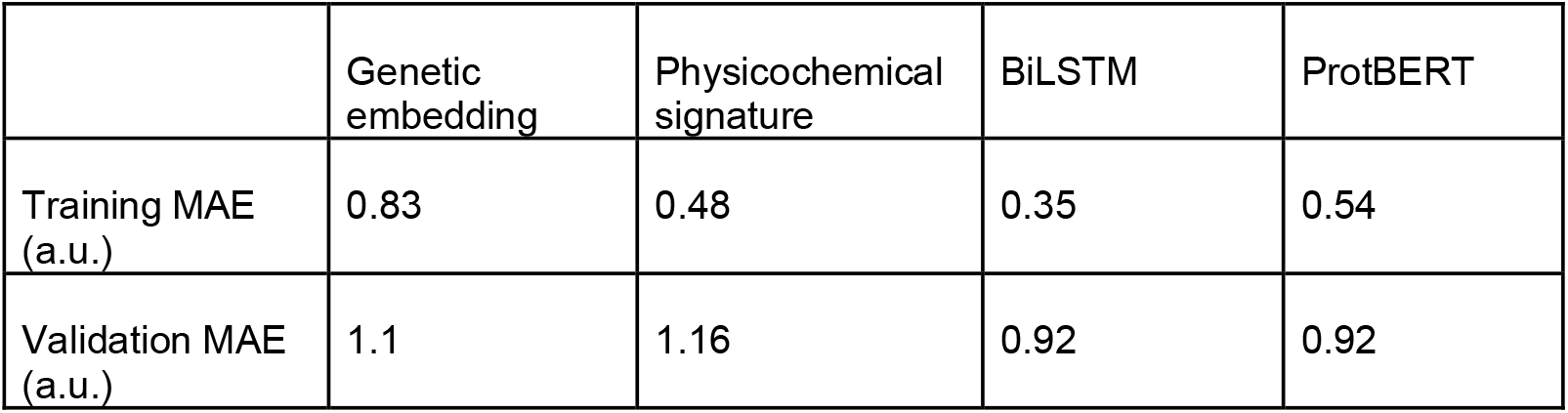
Mean Absolute prediction Error of the LOO-CV ridge regression starting from the embedding vectors of the 4 language models considered.

BiLSTM and ProtBERT embeddings achieved the lowest error on the validation set, even if the overall performances are not strikingly different. For comparison, a null model was generated randomly associating protein sequences and antigenic map coordinates of our training and validation set. The prediction of these random antigenic map coordinates, starting from the 1024-dim vectors of BiLSTM embeddings, has a MAE=3.5 a.u. on the training set and MAE=6 a.u. on the validation set. This shows that the obtained performances on real data are likely not due to an overfitting of a low-dimensional Antigenic Map starting from high-dimensional embeddings. All the embeddings perform well on average, with an error slightly higher than the experimental variability of samples sharing the same HA sequence (0.59 a.u.) and comparable with the experimental error of 0.83 a.u.

In Figure 1 we show the original antigenic map updated until 2011 from Smith et al.^12,15^, while in Figure 2 we show the antigenic maps reconstructed starting from the 4 embeddings. The general outline was similar (see Table 1). Some viruses were not positioned in the correct cluster, depending on the specific embedding type. With the genetic distance embedding, the TX77 cluster was not reconstructed, as it was split between VI75 and BK79 (Figure 2a). For both genetic distance embeddings and physicochemical signatures, the boundary between BE92 and WU95 clusters was not correctly reconstructed (see zoomed regions in Figure 2). The single AA substitution (N145K) causing the difference between viruses of the BE92 and WU95 clusters was captured by both BiLSTM and ProtBERT (Figure 2c-d) and not by the other two embeddings (Figure 2a-b).

**Figure 1.**
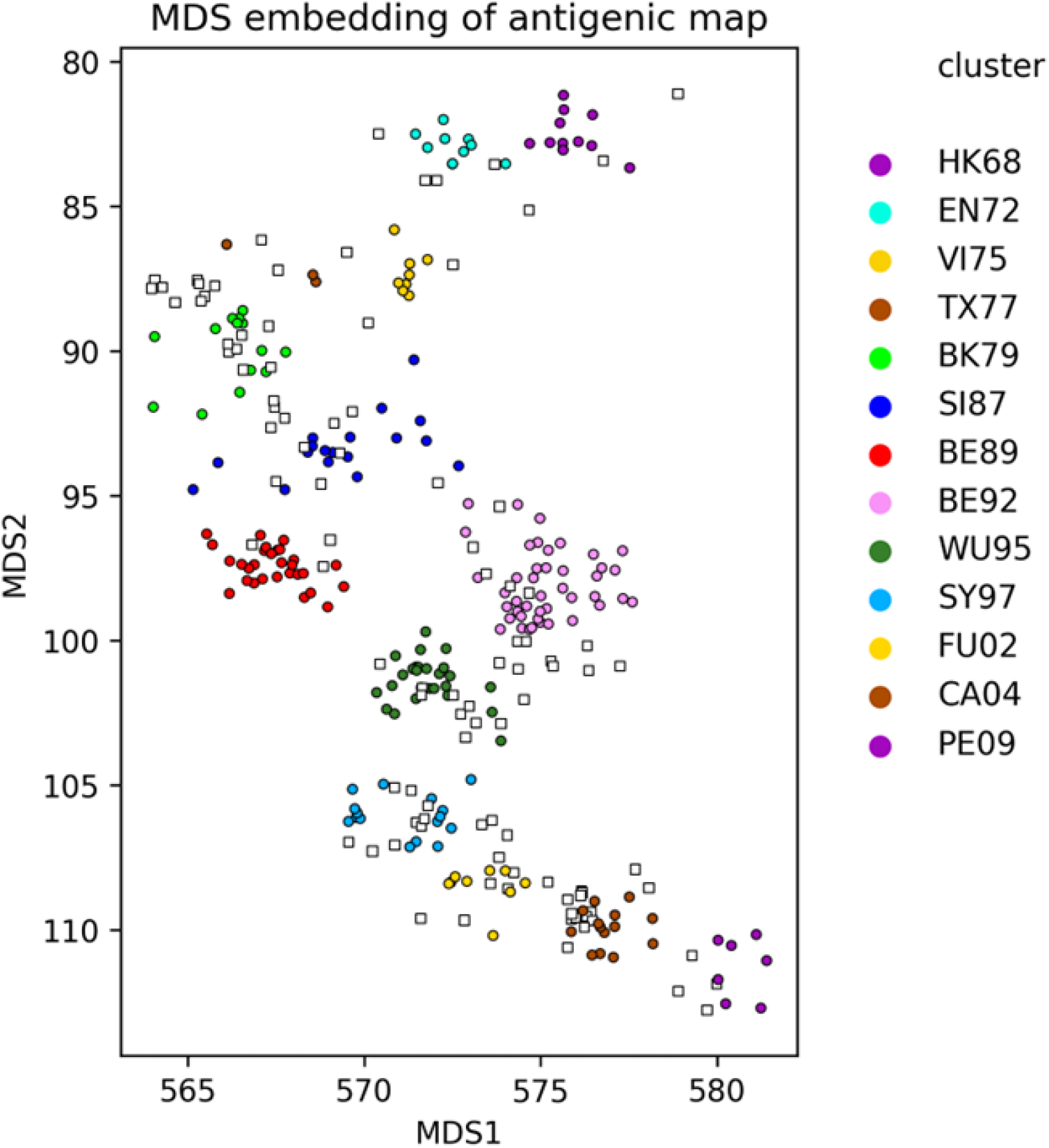
Original antigenic map as in Koel et al.^16^. Antisera are represented as white squares. Antigens are represented as coloured circles, with colors based on the experimental antigenic clusters. Distances between viruses and antisera in the map are inversely related to HI antibody titers.

**Figure 2.**
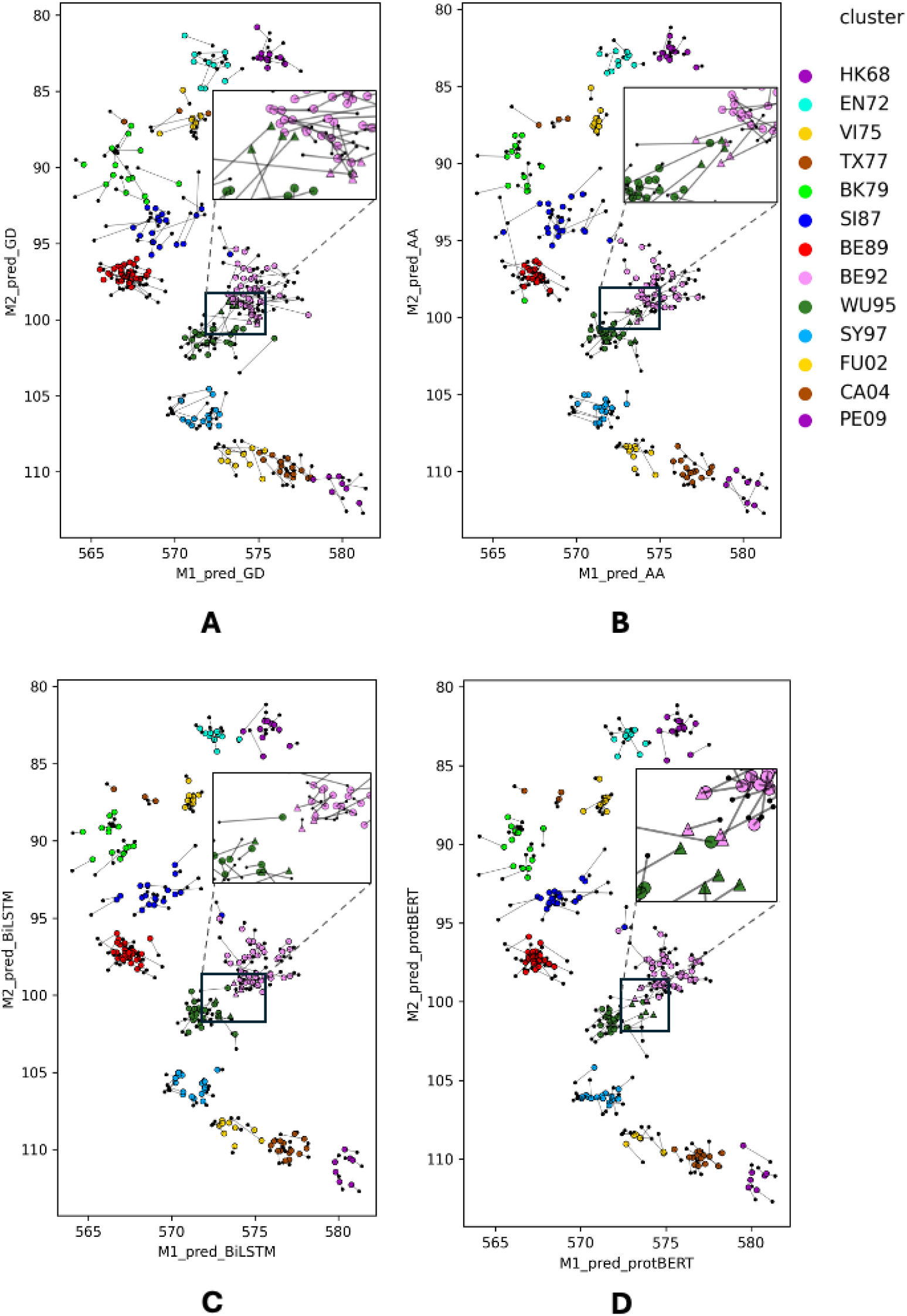
Ridge regression predictions starting from (a) genetic distance embeddings, (b) physicochemical embeddings, (c) BiLSTM embeddings, and (d) ProtBERT embeddings. Black dots represent original experimental AM coordinates, while coloured dots represent the estimated ones, with a line connecting experimental and estimated dots. Viruses of the BE92 and WU95 clusters represented as triangles are of particular interest since they were associated with one cluster in the genetic map based on Hamming distances but in another cluster in the antigenic map as in Smith et al^12^, which was due to a single AA substitution that determines the change in antigenic cluster.

### *in silico* mutational scanning

To evaluate whether our approach can recognize which AA substitutions are causally responsible for major antigenic changes, we generated artificial sequences *in silico* by substituting all the possible AAs in the positions known to cause major antigenic change as described by Koel et al. (145, 155, 156, 158, 159, 189, 193) and predicted their AM coordinates.

The goal was to verify whether the substitutions experimentally associated with antigenic drift (cluster-transition substitutions - CTS) were actually identifiable a-priori with the computational methods presented in this study. The full list of reference sequences and CTS is provided in Supplementary Table S2. We predicted antigenic map coordinates for all the possible single substitutions starting from BiLSTM embeddings, ProtBERT embeddings and physicochemical signatures. This analysis is in fact not applicable to Hamming distance embeddings since all the single substitutions have the same Haming distance from the reference, and thus would have identical embeddings and predicted coordinates.

As an example, in Figure 3 we show for ProtBERT embeddings and physicochemical signatures the predicted positions of mutants starting from the SI87 cluster (Fig. 3a) and starting from BE92 (Fig. 3b), together with the observed samples in that region of the antigenic map. The analogous plot starting from BiLSTM embeddings is shown in Supplementary Figure 1. starting from the virus A/HK/1/89 in the SI87 cluster, represented as a dark red square for the observed position and a dark red cross for the predicted position used to compute the distances. Pink and green dots represent real sequences from two antigenic clusters (BE92 and WU95 respectively) while pink crosses represent mutants starting from virus A/HK/56/94 in the BE92 cluster, represented as a dark red square for the observed position and a dark red cross for the predicted position used to compute the distances. Two multiple substitutions were also applied to the SI87 reference sequence: H155Y_Y159S_R189K and S133D_E156K as these were collectively shown to cause the cluster transitions from the previous and towards the subsequent antigenic clusters^16^.

**Figure 3.**
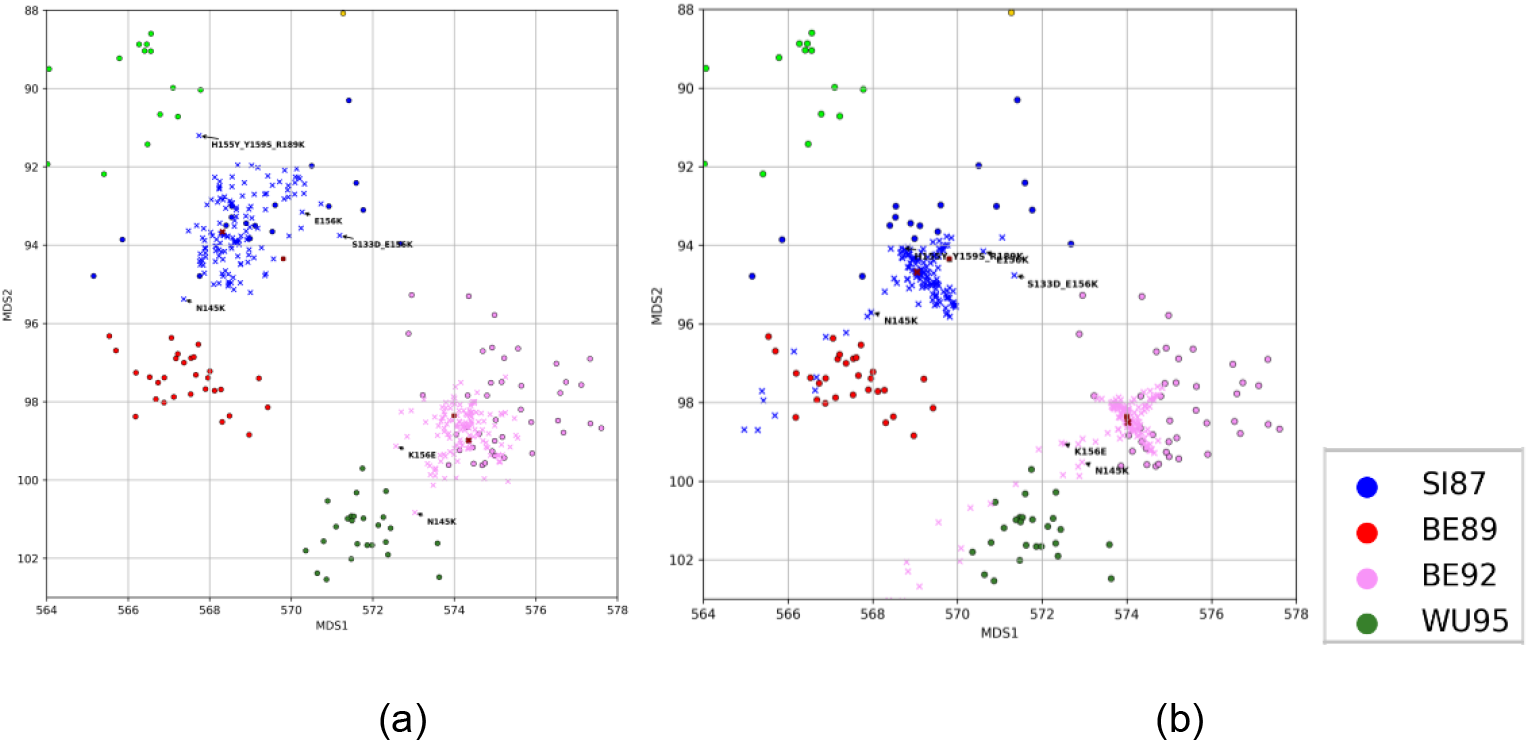
Predicted coordinates for *in-silico* mutants, with a) ProtBERT embeddings and b) physicochemical signatures. Mutant predictions are represented with crosses, observed samples are represented with dots. Blue and red dots represent real sequences from two antigenic clusters (SI87 and BE89 respectively), while blue crosses represent mutants

With ProtBERT and BiLSTM (Figure 3a and Supplementary Fig. S1), the most relevant single CTS for this cluster transition (N145K) moved the mutant sequence in the proximity of the next antigenic cluster (from SI87 to BE89 and from BE92 to WU95), while most of the other mutants remained in the surroundings of the cluster of origin. Importantly, with ProtBERT, N145K was also the substitution with the largest antigenic effect in the correct direction. A similar behavior, but with less magnitude, applies to the other CTS (E156K), which applied to SI87 moves the mutant in the correct direction, which is toward BE92 (see in Figure 1). The same was true in the reverse direction, when K156E was applied to BE92. Given that E156K co-occurred with S133D in SI87, we also tested this combined effect here and this resulted in antigen placement more towards the right direction. It should be noted however that the antigenic distance for all single forward and reverse mutants (E156K, K156E, K145N and N145K) and also for the double mutant was smaller than measured experimentally^16^. We tested a second combination of 3 substitutions, Y155H_S159Y_K189R, known to be responsible for the antigenic change from BK79 to SI87. In our predictions, also this combination mutant correctly moved toward the correct antigenic cluster (i.e. from SI87 to BK79), and the predicted antigenic distance was higher than any single substitution, but still smaller than the experimentally measured distance.

Overall, ProtBERT outperformed the other methods in predicting the antigenic map positions for the in-silico generated single, double and triple mutants, but with an underestimation of the antigenic effect compared to experimental map positions.

Generalizing to all cluster transitions beyond the previous example, the predicted antigenic effect of AA substitutions often did not correspond to the effect of the substitution on antigenic change measured experimentally. Ranking the substitutions based on predicted distance from their reference, the CTS sequences ranked 54/133 for ProtBERT, 46/133 for BiLSTM and 70/133 for physicochemical signatures. This result shows that the distance predicted for single substitutions starting from physicochemical signatures is basically random (70 is approximately mid ranking), while LMs embeddings perform slightly better.

To fully leverage the power of LMs, we decided to utilize also the linguistic probability of each substitution as predicted by the language model, that we call grammaticality as in Hie et al.^8^ (similar to the likelihood that a word fits in a specific position along a sentence). In analogy to what they did in their study, we considered both the distance between sequences and their grammaticality, but our distance was calculated on the predicted 2D antigenic space while the distance used by Hie et al. was calculated in the 1024D embedding space. We thus use a single score we call CAMD (Constrained Antigenic Map Distance score) which is the sum of the rank in antigenic map distance *R*_*D*_ and the rank in grammaticality *R*_*G*_: i.e. a sequence with high CAMD is both likely and distant from the starting point sequence.

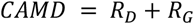

Sorting 133 in-silico substitutions for each of the 13 clusters by CAMD, with equal weight for antigenic distance and grammaticality, and computing the rank of the known CTS, we obtained an average rank of 17/133 for ProtBERT, 21/133 for the BiLSTM (both in the 1st quartile) which improved by far the ranking obtained considering the predicted antigenic distance only (grammaticality score cannot be computed for the other two methods). We also observed that ProtBERT and BiLSTM performances improved if considering only the grammaticality ranking *R*_*G*_ : 5/133 for ProtBERT and 12/133 for BiLSTM. In Supplementary File AM_DMS_rankings_share.xlsx, we list CAMD scores, predicted antigenic distance, and grammaticality for all the *in silico* substitutions. We remark that the *in silico* mutational study was performed starting from 2 different reference sequences in each cluster, for which we obtained consistent results, confirming the robustness of our approach (see also Supplementary Table S3).

### Prediction of future clusters

To investigate if our approaches could identify new antigenic variants that are substantially different from viruses of the current and previous clusters (i.e. variants for which a vaccine update might be needed as it would start a subsequent antigenic cluster), we tested the models through a Leave-the-Future-Out (LFO) validation. To this end, we computed the prediction for samples in the most recent antigenic cluster (PE09) by training the linear RR model only on the previous clusters in time. The LFO approach can be considered equivalent to a real-life situation where a new antigenic cluster has not appeared yet and one tries to place in the map newly observed sequences, even in the case of a single strain, with mutations that have not been observed before. In Table 2 we show the number of PE09 samples correctly recognized as outliers for the CA04 cluster, either according to 95% C.I. or outside a 2 a.u. radius (see also Figure 4).

**Table 2.** Number of PE09 samples (out of 7) correctly recognized as outliers for the CA04 antigenic cluster. First row: samples predicted to be outside a 95% C.I. centered on the cluster centroid. Second row: samples predicted to be outside a 2 a.u. radius centered on the cluster centroid.

| # of PE09 samples recognized as outliers (out of 7) | Genetic distance | Physicochemical signature | BiLSTM | ProtBERT |
| --- | --- | --- | --- | --- |
| 95% C.I. | 2 | 4 | 7 | 6 |
| 2 a.u. radius | 1 | 3 | 5 | 4 |

**Figure 4.**
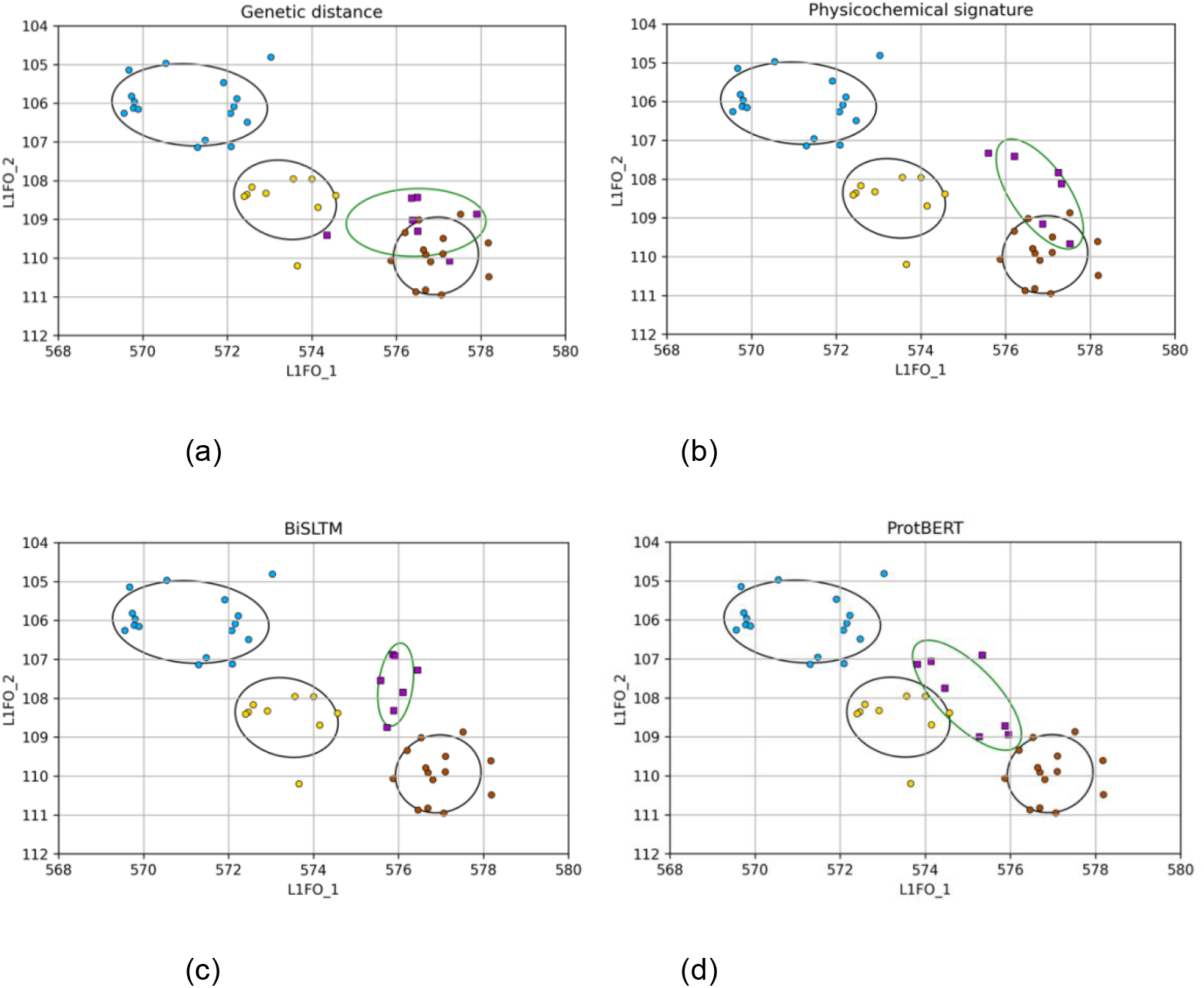
Leave-the-Future-Out prediction for the different used embedding methods (a: Hamming distance, b: physicochemical signature, c: BiLSTM, d: ProtBERT). Dots represent the observed map coordinates of samples used to train the Ridge regression, with the last training set cluster being CA04 (brown dots). Purple squares represent PE09 samples, that have been left out from regression training. Ellipses centered in each cluster’s centroid represent the 68% C.I. of a 2D Gaussian fit.

RR trained on genetic distance embeddings shows a low performance, with only 1-2 held out samples predicted to be outside the most recent training-set cluster (namely CA04, brown dots in Figure 4, and Supplementary File LFO_PE09.xlsx). With physicochemical signatures we correctly recognized only 3 or 4 of the PE09 samples as deviant from CA04. On the contrary, starting from ProtBERT and BiLSTM we recognized as new antigenic variants the majority of held out samples. In particular, with BiLSTM all PE09 viruses were predicted to be outside the 95% C.I. of the previous CA04 cluster. Similarly, BiLSTM and ProtBERT recognized the PE09 cluster to be distinct from CA04 with p-values lower than 0.001 (see Supplementary Table S5). These results showed that Hamming distances and physicochemical embeddings cannot easily quantify the importance of a small number of mutations that lead from an antigenic cluster to the next one, while the other methods are more sensitive to interpreting their effect correctly.

## Discussion

In this study, we compared different embedding methods to represent viral protein sequences, two based on Deep Learning LMs and two based on genetic or physicochemical properties, validating their capability to predict antigenic properties as represented in antigenic maps generated from experimental data. Different from previous approaches aimed to predict antigenic differences from phylogenetic distances^23^ and expert curated features^17–19^, we aimed to start from protein sequences alone for the prediction task. The performance of our models have not been directly compared to existing state-of-the-art models, since they rely on single strain information and not on estimating differences between pairs of strains.

When regressing the embeddings onto antigenic map coordinates, all the approaches achieved a low error on average, comparable to the experimental error. Nonetheless, only Deep Learning language models correctly represented finer-scale features of the antigenic map such as the shape of some antigenic cluster and the shift between antigenic clusters determined by just a single AA substitution.

Through *in silico* mutational scanning we further observed that traditional genetic approaches, such as those based on Hamming distances, do not adequately estimate the impact of specific mutations, quantified by the capability to rank the importance of single AA substitutions correctly. When ranking the most likely substitutions to drive antigenic change, we combined the predicted distance in the antigenic map with the Language Models grammaticality score, that can be interpreted as the likelihood of the mutation to generate a functional sequence, extending the approach proposed in ^8^ by directly mapping the protein LM embeddings into the antigenic space. The CAMD score we defined through the Language Models achieved better results than scores based on genetic distances or physicochemical signatures, although the predicted antigenic distances are often lower than the experimental observations. We also remark that the grammaticality score alone achieved optimal performances, in line with recent studies showing how language models without specific fine tuning can achieve state-of-the-art performance in several deep mutational scanning tasks^24,25^.

Moreover, we performed an ideal experiment mimicking a real-life scenario in which a novel variant emerges, by removing the sequences of the most recent cluster from regression training and calculating their predicted distances with respect to the previous cluster. The Language models, and the physicochemical signatures to a lesser extent, were capable of correctly predicting these sequences as outliers. This result shows the potential of using Deep Learning approaches based on protein sequences alone to identify novel interesting variants before full experimental screening is available, and could possibly alert about the immune escape of these variants. It should be noted that the algorithms correctly identified variants as being antigenically distinct but failed to predict the exact antigenic distance and directionality of antigenic drift. This is true also for experimental data, where new antisera need to be generated against the emerging variants to determine the magnitude and directionality of drift in the map.

In conclusion, Deep Learning LMs perform similarly to the other two methods over coarse features of the antigenic map and significantly better than the other two methods concerning fine-scale properties of the reconstructed antigenic map. We remark that, while BiLSTM was specifically trained on influenza sequences only, ProtBERT had a broader training set built from a wide set of organisms sampled along the full tree of life. It is thus likely that it may show better generalization capabilities also for applications to other viral species, in analogy with LMs applied to human language trained on very broad datasets^26,27^, if provided with sufficient experimental data to reconstruct the evolution of the antigenic landscape.

## Supporting information

LFO_PE09

AM_DMS_rankings_share

## Acknowledgements

This work was supported by the European Union’s Horizon 2020 research and innovation program under grant agreement no. 874735 (VEO) and the National Institute of Allergy and Infectious Diseases–NIH Centers of Excellence for Influenza Research and Response contract 75N93021C00014 (CRIPT).

## Data availability

The code and data used for this study can be accessed on Zenodo with DOI 10.5281/zenodo.14645823. Precomputed embeddings of protein sequences with BiLSTM and ProtBERT can be provided upon request.

## Notes

### Competing Interest Statement

The authors have declared no competing interest.

### Summary of Updates

Minor updates in Results, Figure 2 and Figure 4.

