## Supplementary Materials for "Language models learn to represent antigenic properties of human influenza A(H3) virus"

* corresponding author

### Supplementary Information

#### Supplementary Files

**Suppl. File S1 (a,b).** Results of in-silico mutational scanning. Each sheet contains all the 133 single substitutions we applied to the reference sequence of that cluster, for a) as in Koel et al., for b) with randomly selected reference sequences. is_CTS: 1 if the substitution is a cluster transition substitution. Frequency: observed frequency of that AA in that position in the Smith et al. dataset. Grammaticality (or G) refers to the probability estimated by language models of that AA being observed in that position. Distance (or D) represents the distance in the 2D predicted antigenic map between the mutant and the cluster's reference sequence. CAMD: score computed summing the rank of grammaticality and distance (e.g. ProtBERT_CAMD= Rank_ProtBERT_G + RankProtBERT_D). Mutants are ordered by ProtBERT grammaticality, which resulted the best metric to identify CTS.

**Suppl. File S2.** Leave the Future Out results. For each sequence from the PE09 cluster, we report whether it was recognized as located outside the previous antigenic cluster (CA04) in the map. Each data sheet refers to a different embedding method (ProtBERT, BiLSTM, physicochemical signatures and genetic embeddings).

### Supplementary Tables

| | BiLSTM_128 | BiLSTM_512 | BiLSTM_1024 | ProtBERT |
|---|---|---|---|---|
| Training set LM accuracy | 0.972 | 0.985 | 0.986 | NA |
| AM set LM accuracy | 0.674 +- 0.017 | 0.677 +- 0.02 | 0.894 +- 0.018 | 0.99 +- 0.01 |

**Table S1.** Language Model accuracy (percentage of amino acid correctly guessed, considering all the positions across all the sequences). BiLSTM is trained on 45K HA protein sequences (training set: Hie et al.[1] minus antigenic map sequences from Smith et al.[2]). Both BiLSTM and ProtBERT have also been tested on the 209 protein sequences available for the antigenic maps (AM set). The testing procedure is performed by looping across all amino acids of all protein sequences (e.g. 209x329 loops for the AM set) and predicting the correct amino acid in that position using the rest of the sequence as input. The accuracy is then computed as the fraction of amino acids for which the LM prediction was the same as the observed amino acid.

| **Antigenic cluster: reference sample** | **Cluster transition substitutions (CTS)** |
|---|---|
| HK68: BI/16398/68 | T155Y (F) |
| EN72: BI/23337/72 | Y155T (R), Q189K (F) |
| VI75: BI/1761/76 | K189Q (R), G158E & D193N (F) |
| TX77: BI/2271/76 | E158G & N193D (R), K156E (F) |
| BK79: NL/330/85 | E156K (R), Y155H & S159Y & K189R (F) |
| SI87: SU/1/90 | H155Y & Y159S & R189K (R), N145K (F), E156K (F) |
| BE89: SA/23/92 | K145N (R) |
| BE92: NL/179/93 | K156E (R), N145K (F) |
| WU95: HK/49/95 | K145N (R), K156Q & E158K (F) |
| SY97: NL/427/98 | Q156K & K158E (R), Q156H (F) |
| FU02: NL/222/03 | H156Q (R), K145N & Y159F (F) |
| CA04: NL/348/07 | N145K & F159Y (R), K158N & N189K (F) |
| PE09: PE/16/09 | N158K & K189N (R) |

**Table S2.** Left column: reference sequence of each cluster (from Koel et al.[3]), which means the sequence on which we applied the *in-silico* deep mutational scanning. Right column: list of Cluster Transition Substitutions (CTS), meaning the single substitutions that have been observed to cause a transition to another antigenic cluster when applied to a sequence. Substitutions marked Forward (F) indicate substitutions responsible for the change in antigenic phenotype towards the next antigenic cluster in time, while substitutions marked Reverse (R) indicate substitutions responsible for the change in antigenic phenotype towards the previous antigenic cluster in time. Substitutions joined by the “&” symbol have been shown by Koel et al. to be responsible for antigenic change when applied together.

| | Physicochemical | BiLSTM | ProtBERT |
|---|---|---|---|
| CAMD | 70 +- 37 | 21 +- 22 | 17 +- 19 |
| Grammaticality rank | NA | 12 +- 16 | 5 +- 4 |

(a)

| | Physicochemical | BiLSTM | ProtBERT |
|---|---|---|---|
| CAMD | 60 +- 32 | 30 +- 28 | 17 +- 20 |
| Grammaticality rank | NA | 12 +- 17 | 4 +- 3 |

(b)

**Table S3.** Average rank (out of 133 possible substitutions) of the CTS based on our *in-silico* DMS approach. A lower ranking corresponds to a higher score predicted for the substitution. a) Reference sequences from Supplementary Tab. S2 as a starting point. b) A randomly selected sequence in each cluster is used as a starting point.

| | Phylogenetic | Physicochemical | BiLSTM | ProtBERT |
|---|---|---|---|---|
| Intrinsic dimension | 3.8 | 1.1 | 4.6 | 2.5 |

**Table S4.** Intrinsic dimension of protein embeddings computed with different methods. In previous works[2], [4] the size of antigenic space was estimated between 4 and 5: we estimated the intrinsic dimensionality *ID* for the embeddings obtained with each of the 4 methods following the procedure in [5], which is based on the distance between first and second neighbors.

| | Phylogenetic | Physicochemical | BiLSTM | ProtBERT |
|---|---|---|---|---|
| Probability of PE09 being distinct from CA04 | 0.32 | 0.06 | 0.0006 | 0.00009 |

**Table S5.** Probability of PE09 being distinct from CA04 based on predicted antigenic map coordinates by different embedding methods. Each probability has been estimated computing the Mahalanobis distance of the PE09 predicted centroid from a 2D Gaussian fitted on CA04 samples.

### Supplementary Figures

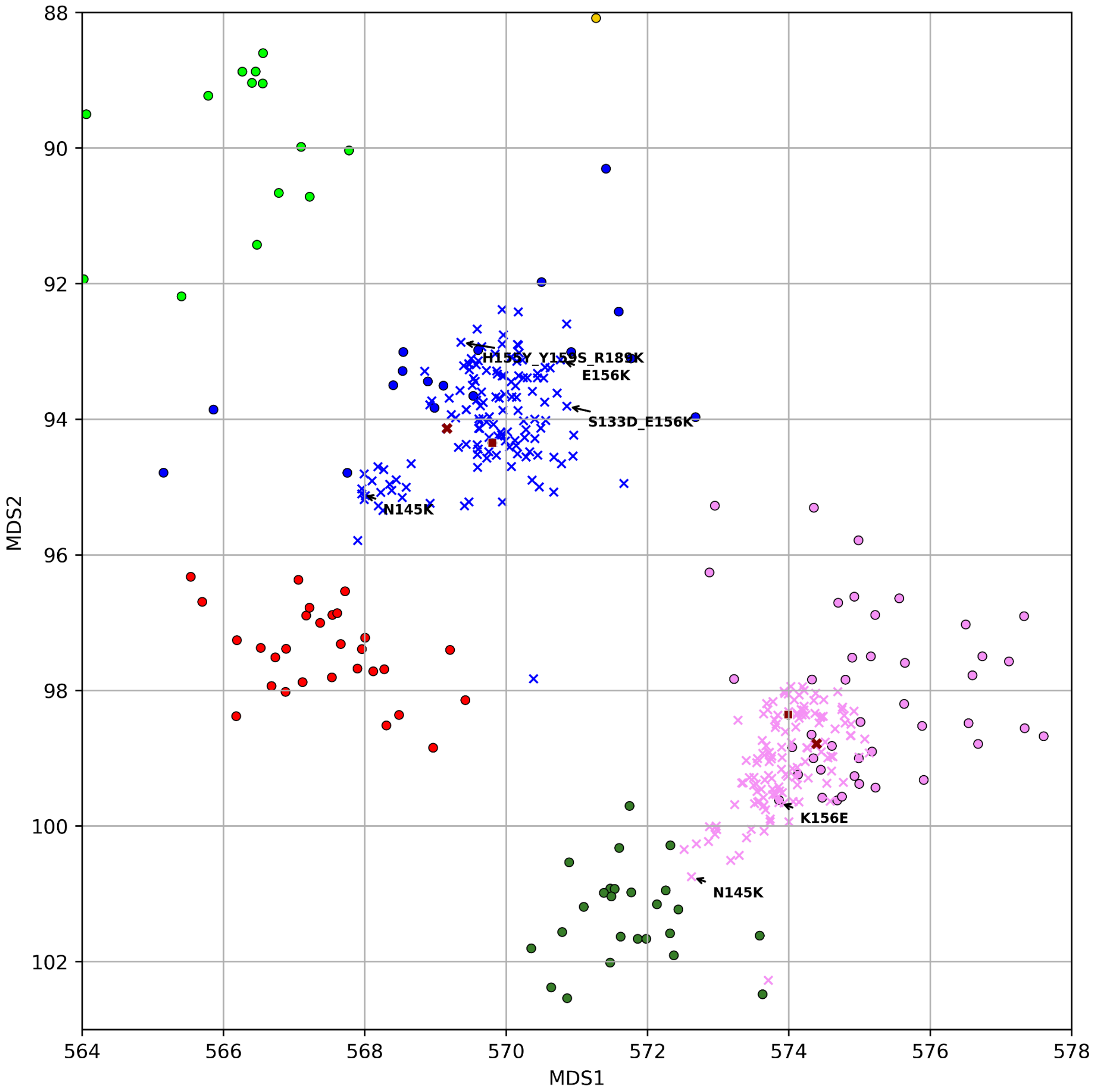


**Figure S1.** Predicted coordinates for single-AA mutants, for BiLSTM embeddings. Mutant sequences are represented with crosses, original experimental samples are represented with circles. Left: pink and green dots represent real sequences from BE92 and WU95 antigenic clusters respectively, while red crosses represent mutants starting from one sequence in the pink cluster (the starting sequence is virus A/HK/56/94 from BE92 cluster, represented as a dark red square for observed coordinates and a dark red cross for predicted coordinates). Right: blue and red dots represent real sequences from BE89 and SI87 antigenic clusters, while blue crosses represent mutants starting from one sequence in the blue cluster (the starting sequence is virus A/HK/1/89 from SI87 cluster, represented as a dark red square for observed coordinates and a dark red cross for predicted coordinates). See also individual mutation rankings in supplementary file AM_DMS_rankings.xlsx.
